# Autocrine feedback maintains homeostatic neuropeptide expression in a peptidergic hub neuron

**DOI:** 10.64898/2026.01.09.698615

**Authors:** Luca Golinelli, Isabel Beets, Liesbet Temmerman

## Abstract

Neuronal circuits sustain stable function by self-regulating their gene expression and neurosecretion. Patterns for autocrine feedback by neuropeptides are widespread across nervous systems, but how they contribute to neuronal homeostasis remains poorly understood. Here, we identify an autocrine homeostatic mechanism in a *C. elegans* peptidergic hub neuron. FLP-1 neuropeptide release activates the inhibitory receptor DMSR-7 on the AVK hub neurons to self-regulate *flp-1* transcription, dense-core vesicle accumulation, and peptide secretion. We show that a parallel PDF-1/PDFR-1 pathway drives *flp-1* expression, establishing a “push-and-pull” peptidergic feedback on the AVK neurons. AVK’s peptidergic output also depends on the number of FLP-1 peptides encoded in its precursor, revealing a dosage mechanism in behavioral regulation. Moreover, *flp-1* expression coordinates transcript levels of additional neuropeptides in AVK. These findings uncover FLP-1 as a self-regulating homeostatic controller within a peptidergic hub neuron and illustrate how precursor-encoded redundancy supports stable yet adaptable circuit states.

## Introduction

Neuronal circuits must balance stability with adaptability, a property maintained through mechanisms of homeostatic regulation ^1–3^. Feedback mechanisms are central to homeostatic control, regulating neuronal excitability in response to activation, and preventing detrimental network states that might lead to hyperexcitability or silencing ^1,4–6^. While synaptic mechanisms of excitation–inhibition balance act on millisecond-to-second timescales, neuropeptides provide an additional regulatory layer that operates over longer durations ^7–9^. Neuropeptides primarily signal by activating G protein-coupled receptors (GPCRs) and can have diverse modulatory effects on neuronal excitability, synaptic efficiency, neurochemical release, and gene expression ^10–14^. Recent advances in single-cell transcriptomic atlases, combined with the large-scale deorphanization of peptide receptors, have enabled systematic reconstruction of neuropeptide signaling networks across species ^15–21^. These neuropeptidergic connectomes reveal a signaling layer that extends beyond canonical wiring diagrams and provides new entry points to dissect the molecular and cellular mechanisms that stabilize circuit dynamics. A recurrent feature of these networks is the high prevalence of possible autocrine feedback loops, in which a single neuron expresses both a neuropeptide and its cognate receptor ^15,17,19^. Such feedback suggests a conserved strategy to restrict the duration of neuropeptide release, sustain activity states over extended periods, or scale neuropeptide production according to demand – mechanisms reminiscent of endocrine feedback regulation ^4,22,23^. Elucidating how feedback mechanisms operate within the broader neuropeptide connectome will be key to understanding their role in circuit stability and behavior.

In *C. elegans*, nearly half of all neurons express cognate neuropeptide receptor-ligand pairs, highlighting this animal’s potential as a model for dissecting peptidergic feedback regulation ^15^. Its compact nervous system, extensively mapped neuropeptidergic connectome, and a comprehensive set of genetic tools and behavioral readouts facilitate targeted studies of these feedback mechanisms ^15,24–26^. Among *C. elegans* neurons, the AVK interneurons are of particular interest for their control of peptidergic output. AVK neurons are peptidergic hub neurons that have been shown to regulate peptide release through autocrine feedback mediated by the RFamide-type neuropeptide FLP-1 ^15,27^. FLP-1 acts via the Gα_i/o_-coupled receptor DMSR-7 to suppress the release of NLP-10 peptides, providing a precedent for hierarchical and homeostatic control within this peptidergic hub ^27^. FLP-1 peptides are primarily released from AVK, can target receptors across the nervous system, and are involved in many AVK-regulated behaviors ^15,27–32^, suggesting widespread regulatory effects. Additionally, the FLP-1 precursor encodes seven highly similar RFamide peptides that activate similar receptors, an arrangement that may have evolved to reinforce and amplify FLP-1 signaling ^20,33^. This raises the question whether and how AVK-derived FLP-1 peptides may act as homeostatic regulators to maintain a stable network.

Here, we show that FLP-1/DMSR-7 signaling acts as a homeostatic control system that represses *flp-1* transcription and peptide release cell-autonomously in AVK neurons. In parallel with DMSR-7 signaling, we identify an excitatory PDF-1/PDFR-1 pathway that positively regulates *flp-1* transcription. Disrupting this equilibrium by removal of FLP-1 peptides reveals redundancy between individual peptide sequences encoded in the *flp-1* gene. Together, these findings support the idea that peptidergic hub neurons in the *C. elegans* nervous system can autonomously adjust their output to maintain behavioral and circuit homeostasis.

## Results

### Inhibitory DMSR-7 signaling in AVK controls *flp-1* expression and behavioral functions

Recent studies highlight a role of FLP-1 neuropeptides in autocrine regulation of AVK neurons. During pathogen avoidance, enhanced AVK activity increases *flp-1* expression, although the underlying mechanisms remain elusive ^29^. In addition, autocrine feedback of FLP-1 on the neuropeptide receptor DMSR-7 was shown to inhibit AVK’s release of NLP-10 peptides ^27^. We sought to determine whether this feedback mechanism acts as a homeostatic controller of AVK’s peptidergic output and investigated whether it self-regulates FLP-1 signaling. To test this, we overexpressed *dmsr-7* in AVK, as it is the only FLP-1 receptor known to be expressed in this neuron class (Figure 1A) ^7,34^, and quantified *flp-1* expression using a CRISPR knock-in transcriptional reporter strain. Overexpression of this Gα_i/o_-coupled receptor significantly reduced AVK fluorescence (Figure 1B-C) ^20,27,29,35^. Besides *flp-1* expression, we also examined whether overexpressing *dmsr-7* leads to impaired locomotor activity, as observed in *flp-1* mutants ^27,28^. In liquid environments lacking food, *C. elegans* engages in accelerated swimming through FLP-1 signaling via the receptors FRPR-7 and NPR-6 ^28^. If increased expression of DMSR-7 in AVK neurons would reduce *flp-1* expression, we could expect to see a corresponding decrease in swimming activity. Indeed, these animals displayed a phenotype similar to *flp-1* null mutants, supporting the idea that DMSR-7 reduces the release of FLP-1 from AVK neurons (Figure 1D). This model is further corroborated by alterations in crawling behavior of transgenic animals. AVK-specific overexpression of *dmsr-7* promoted a faster, roaming-like behavioral state when animals were on food, similar to *flp-1* loss-of-function (Figure 1E-F). We also quantified body angles at five positions along the body axis and found that *dmsr-7* overexpression in AVK enhances body curvature, particularly in the midbody and posterior regions (Figure 1G-H), which is consistent with increased body curvature of *flp-1* mutants ^28,32^. Reintroducing *flp-1* specifically in AVK sufficed to restore both swimming and locomotion on plates (Figure S1A-C).

**Figure 1.**
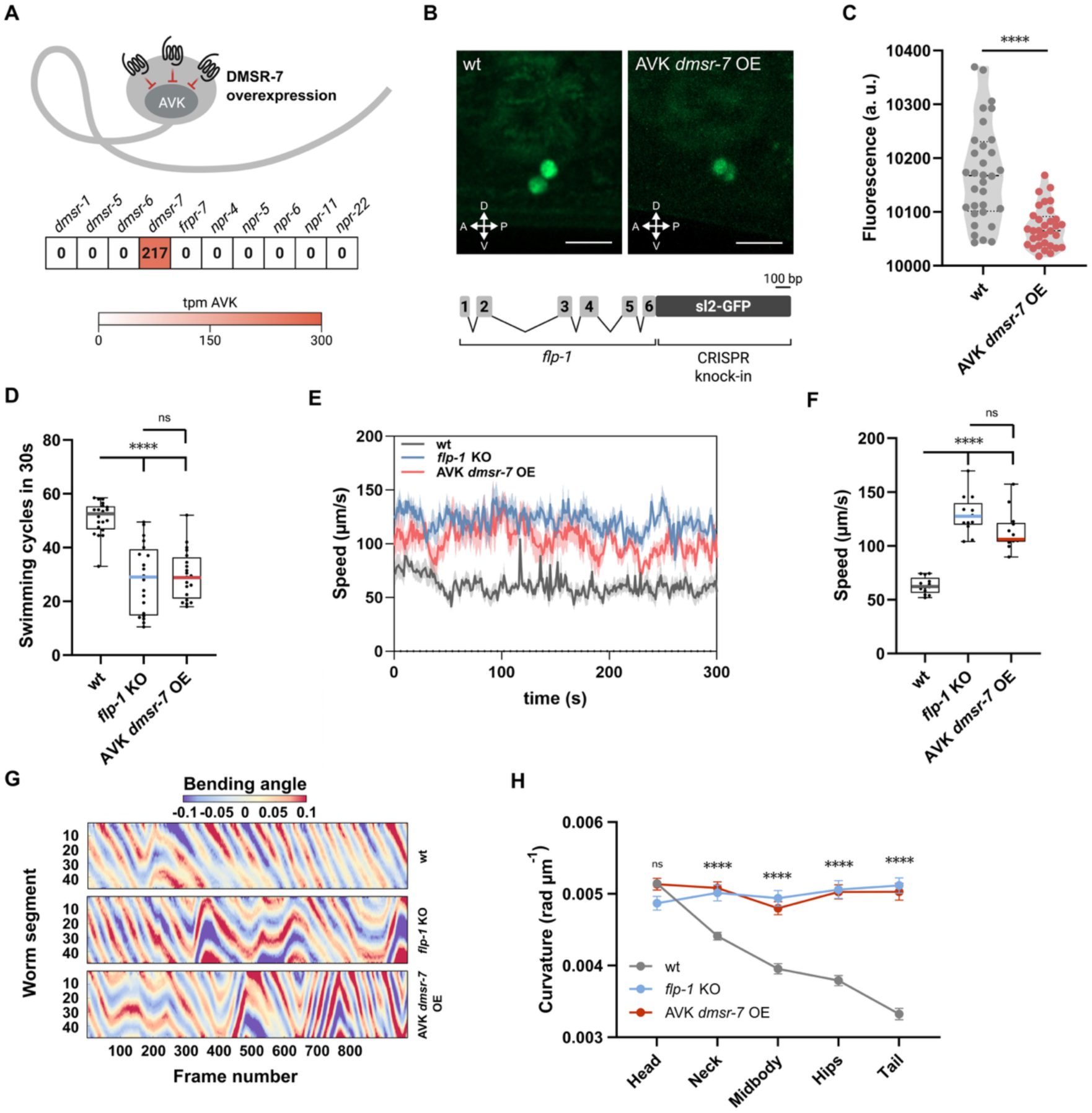
FLP-1 signaling via DMSR-7 in AVK neurons modulates locomotion. **(A)** Expression of *flp-1* receptor transcripts in AVK interneurons, with schematic illustrating targeted overexpression of *dmsr-7* in AVK. **(B)** Confocal image of the *flp-1* transcriptional reporter strain used in this study, showing expression in AVK, alongside a schematic of the CRISPR/Cas9 insertion. Scale bar = 10 µm. **(C)** Quantification of GFP fluorescence intensity in AVK neurons in wild-type and *AVK::dmsr-7* overexpressing animals, indicating successful transgene expression. **(D)** Quantification of total number of swimming cycles in 30 s performed off-food. *flp-1* knockout animals and *AVK::dmsr-7* overexpressing animals display similar reductions in swimming behavior. **(E to F)** Locomotion speed traces **(E)** and quantification of absolute speed on food **(F)** for *flp-1* mutants and *AVK::dmsr-7* animals, showing consistent hyperactivity. **(G)** Kymographs illustrating body curvature across the anterior–posterior axis in freely moving animals. **(H)** Quantification of mean body curvature showing increased bending in both *flp-1* mutants *and AVK::dmsr-7* animals. Data show mean ± SEM from N ≥ 14 per strain. Statistical comparison among strains was performed using t-test **(C)**, one-way ANOVA with post hoc Dunnet’s test **(D)** and Tukey test **(F and H)**; ****p < 0.0001

To further support the hypothesis that DMSR-7 signaling regulates FLP-1 synthesis, we examined if loss of *dmsr-7* alters *flp-1* expression. We first extracted RNA from whole animals and compared *flp-1* transcript levels between wild-type and *dmsr-7* mutant animals. As expected, *flp-1* transcripts were significantly increased in *dmsr-7* null mutants (Figure 2A). To validate these findings at the cellular level, we compared fluorescence of the *flp-1* transcriptional reporter in AVK between wild type and *dmsr-7* mutants. Disrupting *dmsr-7* resulted in increased AVK fluorescence, indicative of elevated *flp-1* expression (Figure 2B). To rule out the possibility that this effect was mediated by other neurons expressing DMSR-7, we generated an AVK-specific *dmsr-7* knockout line using Cre recombinase. This targeted deletion phenocopied the increased expression of *flp-1* observed in the *dmsr-7* null mutant, confirming that DMSR-7 cell-autonomously regulates *flp-1* expression in AVK (Figure 2B).

**Figure 2.**
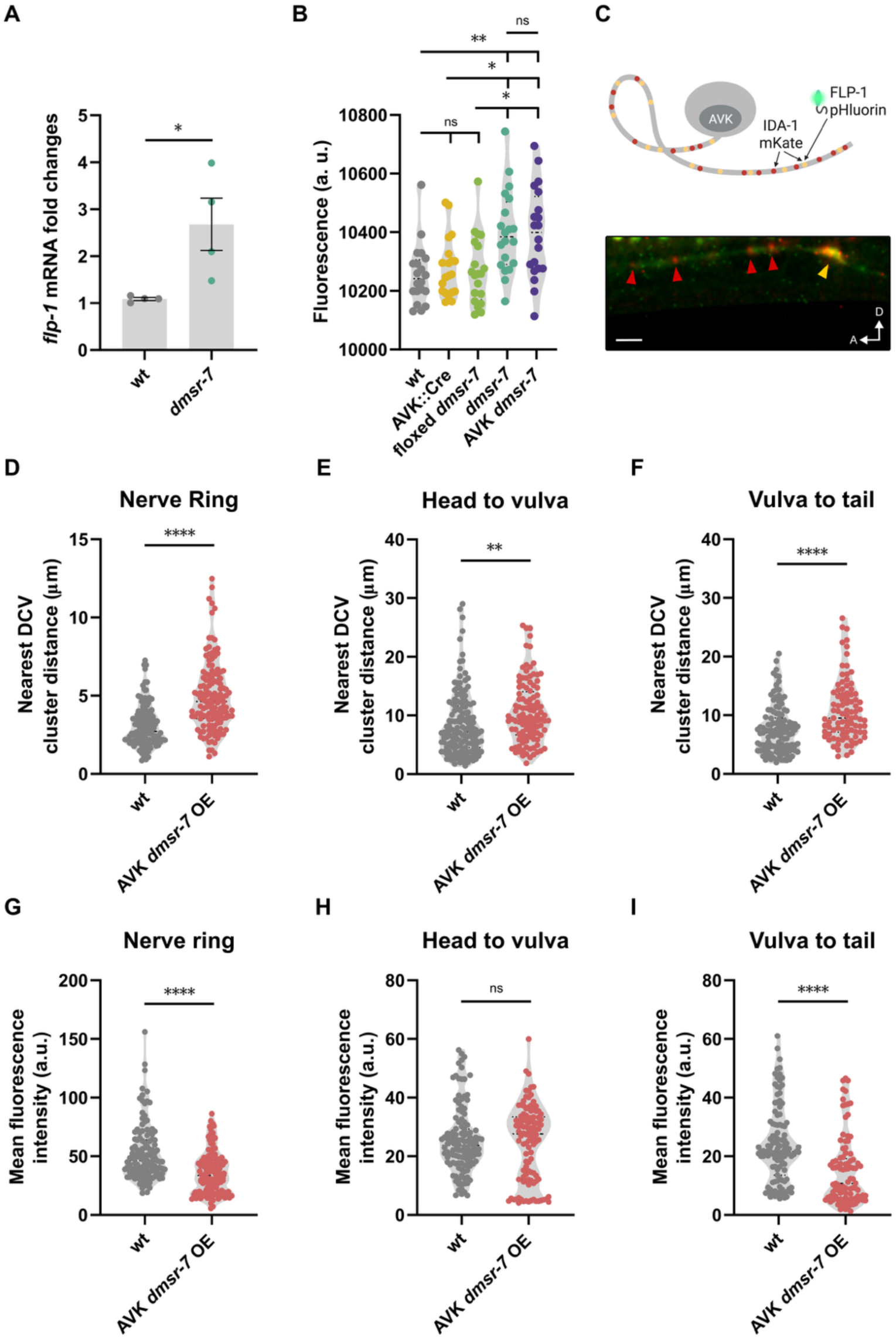
Loss of DMSR-7 increases *flp-1* expression and alters dense core vesicle distribution. **(A)** Relative expression levels of *flp-1* mRNA in whole-animal extracts from wild-type and *dmsr-7* knockout animals, showing a significant increase in *flp-1* transcript abundance in the absence of *dmsr-7*. **(B)** Mean AVK fluorescence intensity using a *flp-1* transcriptional reporter strain, showing elevated expression in both whole-animal *dmsr-7* knockouts and AVK-specific *dmsr-7* deletion strains, compared to wild type animals. **(C)** Schematic and representative confocal image of the dual-reporter strain used to assess *flp-1*-containing dense core vesicles (DCVs). Green puncta indicate *flp-1::pHluorin*; red puncta mark *ida-1::mKate*-labeled DCVs; orange puncta indicate the overlap of the two. Scale bar = 5 µm. **(D to F)** Quantification of inter-puncta distances for *flp-1*-positive DCVs in three anatomical regions in non-transgenic *unc-32* mutants and *unc-32* mutants overexpressing *dmsr-7* in AVK: within the nerve ring **(D)**, in axon regions extending from the neck to the vulva **(E)**, and from the vulva to the tail **(F)**. **(G–I)** Mean fluorescence intensity of *flp-1::pHluorin*-positive DCVs in non-transgenic *unc-32* mutants and *unc-32* mutants overexpressing *dmsr-7* in AVK: in the nerve ring region **(G)**, in axons sections from neck-to-vulva **(H)**, and vulva-to-tail tip axons (**I**). All data represent mean ± SEM. **(A)** N = 3 independent assays; **(B) and (D to I)** N ≥ 20 animals per genotype. Statistical analysis performed using t-test **(A and D-I)** one-way ANOVA with post hoc Tukey correction **(B)**; ****p < 0.0001, ***p < 0.001, **p < 0.01, *p < 0.05.

### DMSR-7 regulates dense-core vesicle accumulation and FLP-1 release in AVK neurons

We next asked whether FLP-1/DMSR-7 signaling regulates the release of FLP-1 peptides, in addition to *flp-1* gene transcription, in AVK. Unlike classical neurotransmitters, which are stored in small synaptic vesicles and released at presynaptic terminals, neuropeptides are packaged into dense-core vesicles (DCVs) that can be released from varicosities or distal neuronal processes ^36,37^. We investigated if FLP-1/DMSR-7 signaling affects the distribution of DCVs in AVK by tagging FLP-1 with pHluorin, a pH-sensitive green fluorescent protein that is quenched in the acidic environment of DCVs (pH < 5.5) ^38^. To visualize the tagged peptide, we crossed this strain with an *unc-32* mutant, which lacks a functional subunit of the vesicular proton pump, thereby increasing intravesicular pH and unquenching the pHluorin signal ^39^. To mark vesicles within AVK, we co-expressed a fluorescently tagged version of IDA-1, a protein associated with both DCV release and vesicle recycling ^38^. Due to the involvement of IDA-1 in both processes ^38^, we observed more IDA-1-positive puncta than pHluorin-positive puncta in AVK (Figure 2C). In control animals, most puncta were clustered in the nerve ring and more dispersed vesicles appeared along neuronal processes (Figure 2C-I). By contrast, vesicles appeared to be more spaced in transgenic animals overexpressing *dmsr-7* in AVK (Figure 2D-F), and fluorescence intensity was reduced in both the nerve ring and AVK’s processes (Figure 2G-I). This suggests that DMSR-7 controls both FLP-1 synthesis and DCV organization in AVK.

To probe if DMSR-7 regulates FLP-1 secretion, we monitored coelomocyte uptake of mKate2-tagged FLP-1 peptides ^27^. Animals overexpressing *dmsr-7* in AVK showed reduced fluorescence in coelomocytes in comparison to controls animals, indicative of decreased peptide release (Figure S1D). We further tested this hypothesis using a nanotrap-based approach^41^ to capture secreted FLP-1::pHluorin peptides with a GFP-binding protein (GBP) fused to the membrane-anchored protein SAX-7 ^39–41^. We expressed the GBP::SAX-7 fusion protein under the control of the cholinergic *unc-17* promoter (Figure S1E). Notably, *unc-17* is expressed in several FLP-1 target neurons, including AIM, SMB, and VC, but not in AVK, minimizing peptide trapping at the site of release ^28,32,34^. In transgenic animals co-expressing *flp-1::pHluorin* and *unc-17p::GBP::sax-7*, we observed numerous fluorescent neurons predominantly in the head region as well as along the ventral nerve cord and vulval region, confirming a broad signaling range of FLP-1 (Figure S1G–H). Interestingly, overexpression of *dmsr-7* diminished the number of fluorescent neurons (Figure S1F-H), suggesting that FLP-1 release is strongly reduced in these animals.

### Homeostatic regulation of *flp-1* expression in AVK occurs via autocrine FLP-1/DMSR-7 feedback

DMSR-7 is a promiscuous receptor that, in cell-based receptor activation assays, can be activated by FMRFamide-like peptides encoded by 29 *flp* precursor genes in the *C. elegans* genome ^20,42^. Because of this promiscuity, FLP-1 regulation in AVK may result from autocrine feedback of FLP-1 or from FLP signaling mediated by other peptide sources. To determine if AVK-derived FLP-1 could self-regulate its transcription, we used a biochemical approach to tether FLP-1 peptides to the surface of AVK and quantified *flp-1* expression (Figure 3A) ^43,44^. We hypothesized that this physical constraint elicits constant activation of DMSR-7 on AVK, overactivating its inhibitory intracellular cascade. As expected, we observed a decrease in *flp-1* transcription in AVK when tethering the peptide to AVK’s surface (Figure 3B). This result suggests that constant FLP-1 signaling in AVK diminishes the production of FLP-1 peptides in this neuron. Indeed, animals expressing the AVK-tethered FLP-1 transgene show behavioral defects in both swimming and crawling on plates, reminiscent of *flp-1* null mutants (Figure 3C-E). As a control, we also tethered another peptide, NLP-1, to AVK’s surface. NLP-1 does not bind to DMSR-7, and its cognate receptors (NPR-9 and NPR-11) are not expressed in AVK ^34,45,46^. Tethering NLP-1 on AVK’s surface, unlike FLP-1, did not reduce peptide expression in comparison to control animals, and animals expressing the AVK-tethered NLP-1 transgene showed normal behavior (Figure 3B-E). Taken together, these results suggest that FLP-1 can interact with receptors on the surface of AVK to suppress its own transcription.

**Figure 3.**
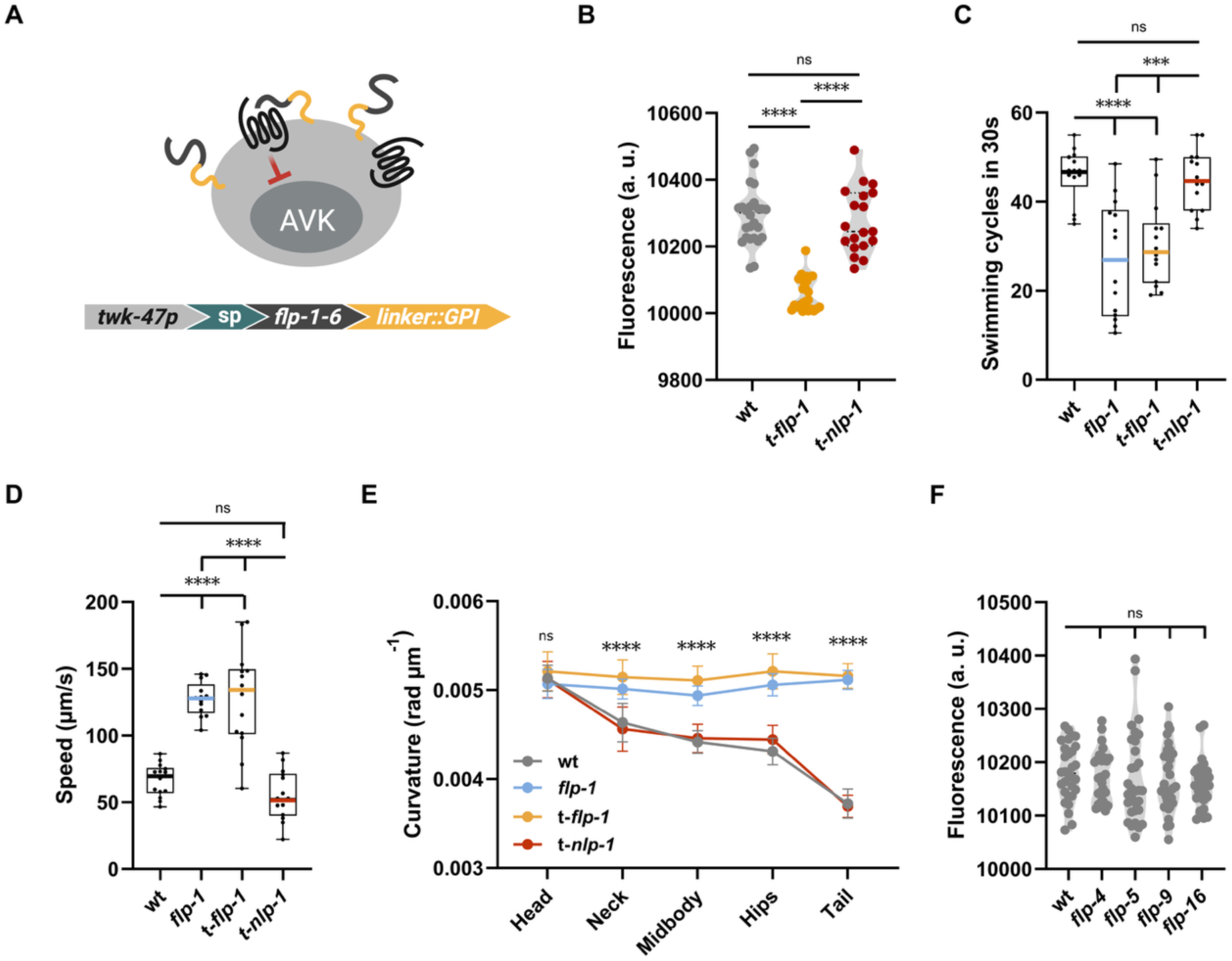
AVK derived FLP-1 and PDF-1 signaling converge on *flp-1* regulation and behavioral output. **(A)** Schematic of the biochemical strategy used to tether neuropeptides to the membrane of AVK neurons. Constructs included the AVK-specific *twk-47* promoter, a signal peptide (sp), either *flp-1-6* or *nlp-1-1* neuropeptide sequences, and a glycosylphosphatidylinositol (GPI) anchor to localize peptides to the neuronal surface. **(B)** Quantification of mean AVK fluorescence using a *flp-1* transcriptional reporter strain shows increased expression in animals expressing tethered *flp-1* (t-*flp-1*) or *nlp-1* (t-*nlp-1*) in AVK, compared to wild type. **(C)** Swimming cycles during a 30 s off-food assay. Animals expressing t-*flp-1* in AVK recapitulate the behavioral phenotype of *flp-1* knockout mutants. **(D)** Mean body curvature measured at three body segments, revealing increased curvature in *flp-1* mutants and AVK t-*flp-1* animals. **(E)** Mean crawling speed on food, showing reduced locomotion in *flp-1* KO and t-*flp-1* animals. **(F)** AVK fluorescence in *flp-1* reporter animals lacking high-affinity DMSR-7 ligands shows no change, suggesting these ligands do not regulate *flp-1* expression in AVK. All fluorescence data represent mean ± SEM from ≥ 20 animals per genotype; behavioral data represent mean ± SEM from ≥ 14 independent assays. Statistical analysis by one-way ANOVA with post hoc Dunnett’s test **(C)**, and Tukey test **(B, D, E, and F)**; ****p < 0.0001, ***p < 0. 001.

One possibility is that downstream target neurons provide feedback to AVK to suppress FLP-1 release, a common feedback motif in endocrine signaling cascades. Previous studies have identified AIM, NSM, SMB, and VC neurons as FLP-1 targets in the regulation of locomotion and body posture ^28,32^. Collectively, these neurons express 18 FMRFamide neuropeptides, with FLP-4, FLP-5, FLP-9, and FLP-16 being the most potent ligands of DMSR-7 alongside FLP-1 ^20,34^. Among these, *flp-16* is the only other *flp* gene expressed in AVK, potentially mediating autocrine feedback via DMSR-7 ^34^. To probe if these peptides regulate *flp-1* transcription in AVK, we quantified *flp-1* expression in loss-of-function mutants. We observed no change in AVK fluorescence, suggesting that these peptides are not individually able to regulate *flp-1* expression (Figure 3F). Collectively, these findings support a model where FLP-1 self-regulates its expression by autocrine feedback through DMSR-7 in AVK neurons. They also show that other high-potency receptor ligands do not significantly influence *flp-1* expression, although we cannot rule out the possible involvement of FLP peptides not included in our screening.

### FLP-1/DMSR-7 and PDF-1/PDFR-1 signaling antagonistically regulate neuropeptide expression in AVK

Our analysis into the potential roles of other neuropeptides regulating *flp-1* expression did not reveal any significant changes. We next reversed our focus to investigate whether FLP-1 might, in turn, influence the expression of other peptide-encoding genes—particularly those expressed in AVK. Transcriptomic data show 14 additional neuropeptides expressed in this peptidergic hub neuron (Figure S2A-B) ^15,25,34^. Among these, the release of NLP-10 peptides was shown previously to be regulated by FLP-1/DMSR-7 signaling, although *nlp-10* expression is not affected in *flp-1* mutants ^27^. We asked whether loss of *flp-1* might affect the expression of other AVK-derived neuropeptides. To probe this, we extracted RNA from whole mount wild type and *flp-1* mutant animals and compared the transcript levels of 14 AVK-expressed neuropeptides among the two strains. We found three neuropeptide genes—*nlp-21*, *nlp-49* and *nlp-50*—to be differentially expressed in *flp-1* mutants. For *nlp-49* we observed reduced expression, whereas transcript levels of *nlp-21* and *nlp-50* were increased (Figure S2C). Since these neuropeptide genes are expressed in multiple neurons, in addition to AVK, altered neuropeptide expression in whole-mount RNA samples may reflect alterations in different neurons. To test if *flp-1* regulates peptide expression in AVK, we compared the expression levels of *nlp-49* and *nlp-50* in wild-type and *flp-1* mutant animals using CRISPR knock-in transcriptional reporters. We crossed these reporter strains with a fluorescent landmark strain carrying the NeuroPAL transgene to facilitate AVK identification ^15,47^. Quantification of GFP fluorescence in AVK confirmed that *nlp-49* expression is reduced in *flp-1* mutants, while expression of *nlp-50* is increased (Figure 4A-B).

**Figure 4.**
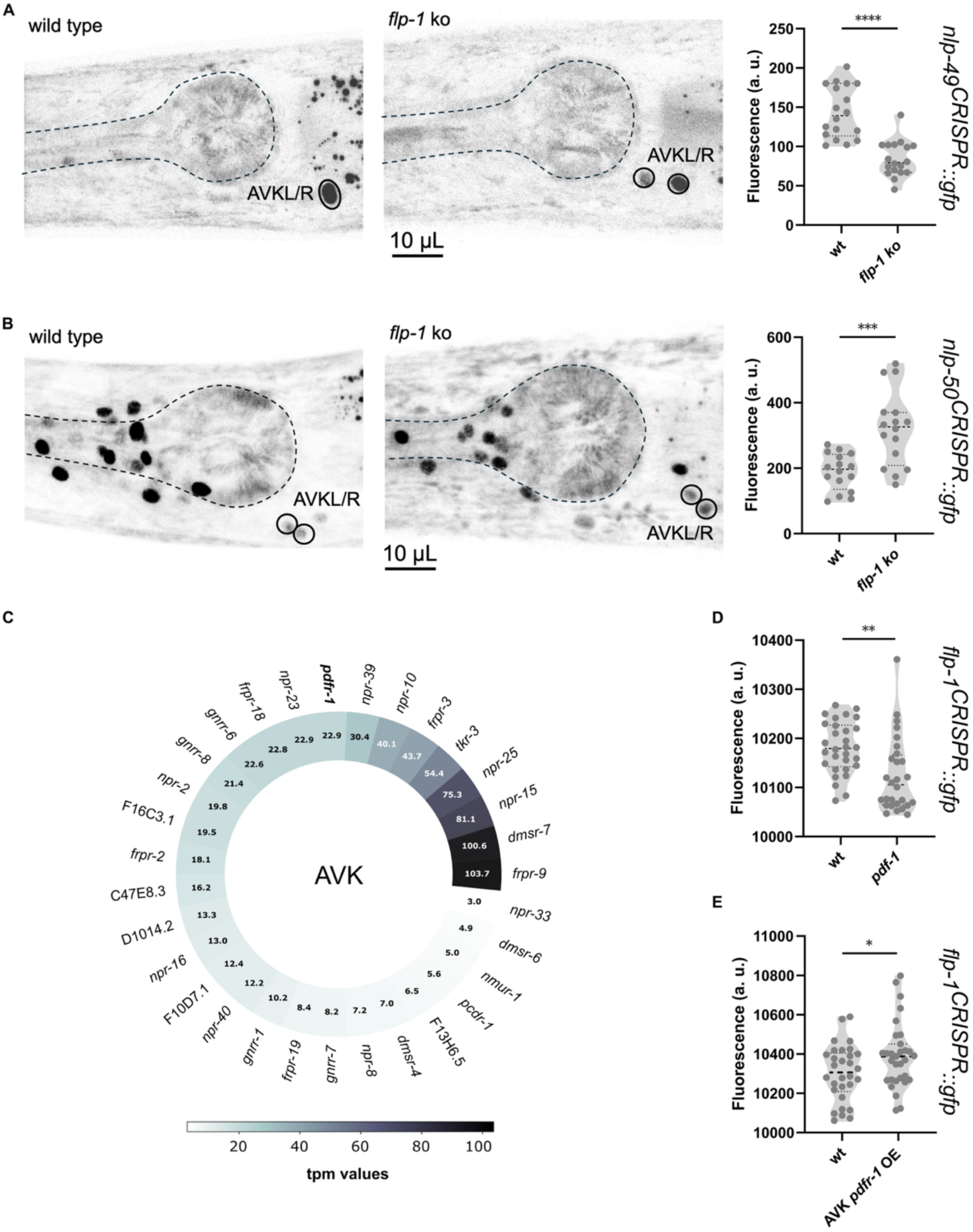
FLP-1 and PDFR-1 influence the peptidergic profile of AVK. **(A)** Confocal micrographs and quantification of *nlp-49* reporter fluorescence in AVK nuclei of wild-type and *flp-1* mutants. **(B)** Confocal micrographs and quantification of *nlp-50* reporter fluorescence in AVK nuclei of wild-type and *flp-1* mutants. **(C)** Expression of neuropeptide receptors in AVK neurons according to CeNGEN, threshold 2 ^34^. **(D)**, *pdf-1* mutants exhibit significantly reduced *flp-1* expression, indicating a role for PDF-1 in regulating *flp-1* levels. **(E)**, Overexpression of *pdfr-1* in AVK neurons increases *flp-1* transcriptional reporter signal, supporting a cell-autonomous role for PDF-1 signaling. All fluorescence data represent mean ± SEM from ≥ 15 animals per genotype. Statistical analysis by t-test; ****p < 0.0001, ***p < 0.001, **p < 0.01, *p < 0.05.

Our findings show that FLP-1 feedback on AVK neurons not only self-regulates *flp-1* expression but also controls transcription of other AVK-derived peptides. We next asked whether inhibitory feedback through FLP-1/DMSR-7 signaling is antagonized by an excitatory pathway, which may increase peptide expression in AVK neurons. AVK expresses 31 neuropeptide receptors (Figure 4C), among which PDFR-1 is the only GPCR known to couple with a Gαs protein ^48^. Notably, AVK also produces the neuropeptide PDF-1, the cognate ligand of PDFR-1 ^34^. Because DMSR-7 and PDFR-1 signal through G protein pathways with opposing effects, we asked if PDF-1/PDFR-1 signaling also contributes to the regulation of *flp-1* expression. We first examined *flp-1* transcript levels in a *pdf-1* mutant and found that *flp-1* expression is reduced, consistent with the excitatory role of PDFR-1 (Figure 4D) ^48,49^. Next, we investigated if PDFR-1 signaling promotes *flp-1* expression by overexpressing the receptor specifically in AVK. Increased *pdfr-1* expression enhanced the expression of *flp-1,* opposite to the effect of *pdf-1* loss-of-function (Figure 4E). These results show that PDF-1/PDFR-1 signaling regulates *flp-*1 expression antagonistically to FLP-1/DMSR-7 in AVK neurons.

### FLP-1 regulates locomotion in a copy number-dependent manner

Our findings indicate that maintaining appropriate FLP-1 levels is essential for normal locomotion. These levels are controlled by a negative autocrine feedback loop involving DMSR-7, counteracted by PDFR-1 signaling in AVK. Since the FLP-1 precursor encodes seven RFamide peptides that can activate similar receptors (Figure S3A) ^20^, we wondered if these copies might act redundantly in the regulation of *flp-1* expression and release. To test this, we genetically removed copies of the RFamide peptides from the *flp-1* gene while preserving the overall gene structure and precursor’s backbone (Figure 5A). We found that *flp-1* mRNA levels increase proportionally to the number of FLP-1 peptides removed from the precursor. The strongest increase occurred in strains that retain none (without RYamide and RFamide peptides) or only a single copy of FLP-1 peptides. By contrast, *flp-1* expression was normal in strains where three or more RFamide peptides were preserved in the FLP-1 precursor (Figure 5B), suggesting there might be some functional redundancy between individual FLP-1 peptides. To further test this hypothesis, we assayed the animals in liquid and scored their swimming cycles. Consistent with our expression data, three copies of RFamide-type peptides in the FLP-1 precursor sufficed to restore swimming to wild type levels (Figure 5C). To further explore the relationship between peptide copy number and behavior, we also examined the speed and body curvature of these animals when crawling on a bacterial lawn. In this case, two copies of FLP-1 RFamide peptides sufficed to restore normal locomotion (Figure 5D-E).

**Figure 5.**
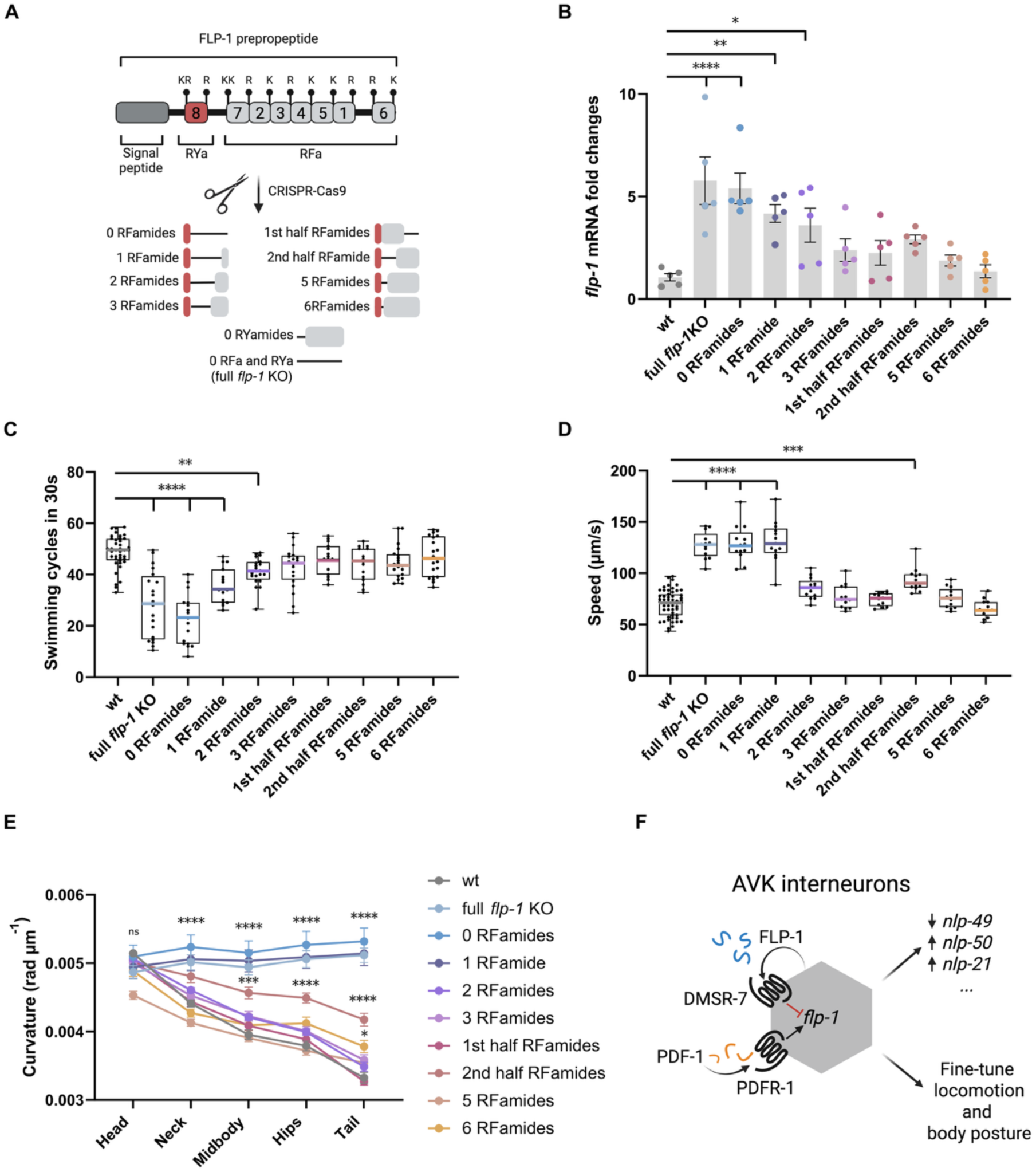
FLP-1 RFamides dosage shapes AVK-driven motor control through compensatory regulation. **(A)**, Schematic of the CRISPR/Cas9 genome editing strategy used to generate a series of *flp-1* alleles with reduced numbers of RFamide and RYamide-encoding peptides. Details in Supplementary file S1. **(B)**, Quantification of *flp-1* mRNA levels in the generated strains reveals compensatory upregulation in animals lacking all or most FLP-1 RFamides. **(C)** Swimming cycles performed during a 30 s off-food assay, showing progressive behavioral deficits correlating with decreasing neuropeptide copy number. **(D and E)**, Mean locomotion speed **(D)**, and body curvature **(E)** measurements of *flp-1* copy number mutants, indicating altered movement dynamics and postural control. **(F)**, working model: AVK neurons create an autocrine loop via FLP-1/DMSR-7 interaction to fine-tune their own neuropeptide output and regulate locomotion and posture. This local feedback mechanism acts hierarchically to modulate the expression of other neuropeptides contributing to motor behavior. PDF-1 from other neuronal sources provides an external modulatory input to balance AVK activity. All data represent mean ± SEM from ≥ 14 independent assays per genotype. Statistical analysis performed using one-way ANOVA; ****p < 0.001, ***p < 0.001, **p < 0.01, *p < 0.05.

Besides RFamide-type peptides, the FLP-1 precursor also encodes a single copy of an RYamide peptide (Figure S3A-B), which can activate DMSR-7 in heterologous cells ^20^. We therefore asked whether this peptide also self-regulates *flp-1* expression. However, we observed no alterations in behavior or *flp-1* expression in a strain lacking the RYamide-type peptide (Figure S3C-F), with only a mild increase of hips and tail curvature (Figure S3F). Expression of *flp-1* also did not change when the RYamide peptide was removed from a strain expressing only three RFamides (Figure S3C), further corroborating that there is no additive effect on expression. Taken together, these data suggest that the FLP-1 precursor encodes more RFamide peptides than strictly needed for normal locomotion, potentially maintaining the capacity to concomitantly regulate other functions. Alternatively, additional copies may be essential under non-standard conditions, such as food deprivation or exposure to noxious stimuli, where sustained release of AVK-derived FLP-1 could be necessary.

## Discussion

Neuropeptidergic autocrine feedback loops are widespread across species, suggesting they likely emerged as an early strategy to control neural signaling dynamics ^1,6,15,17^. Examples of this form of neural regulation have been studied in several phyla, such as the *Aplysia* neurotrophin ApNT, which activates ApTrk to enhance glutamate release ^50^, and the cross-regulation of vasopressin and dynorphin in mammals ^51^. Peptidergic feedback has been shown in *C. elegans* as well, for example, in BAG neurons, where insulin expression is repressed by autocrine feedback of insulin-like peptides during neuron development ^11^. Similarly, the autocrine interaction between FLP-11/DMSR-1 in the RIS neuron regulates the duration of sleep ^52^, and FLP-1/DMSR-7 signaling in AVK controls NLP-10 release ^27^. Our results extend these examples and identify a peptidergic self-regulatory mechanism in AVK that sustains homeostatic control of neuropeptidergic output. This mechanism may be a recurring feature of peptidergic hubs, as suggested by neuropeptidergic connectome data, enabling neurons to scale peptide synthesis and release according to environmental conditions and biological demands ^15,17^.

To counterbalance the inhibiting effect of FLP-1/DMSR-7 signaling in AVK, we show an excitatory PDFR-1 signaling pathway, which boosts *flp-1* expression. Such a push-and-pull mechanism resembles classical endocrine loops, such as the GnRH-LH/FSH axis, where stimulatory signals are paralleled by inhibitory feedback mechanisms to maintain proper balance ^53^. It remains unclear whether FLP-1 also influences PDF-1 signaling within AVK or other neuronal sources, but reciprocal regulation would provide a fine-tuning mechanism for locomotor output. A similar role of PDF signaling in sustaining peptidergic release has been described previously in *Drosophila* pacemaker cells, where PDF engages in autocrine loops to promote persistence of the circadian state ^43^. Thus, AVK likely integrates opposing peptidergic pathways to control behavioral output, highlighting hub neurons as dynamic nodes with the potential to shift state depending on physiological context (Figure 5F).

A further insight is that copy number of FLP-1 RFamide peptides influences AVK-dependent behaviors. We find that three or more FLP-1 RFamide copies suffice to mediate normal locomotion, indicating a dose-dependent action of FLP-1 peptides. Similar results were reported in knockout strains rescued with truncated versions of the gene, which encode only a subset of the peptides ^27^. This suggests functional redundancy between multiple peptides encoded in the FLP-1 precursor. Nematode RFamide peptide precursors often encode two or more copies, while some mollusks and arthropods can have over ten ^42,54–61^. This organizational pattern is also evolutionarily conserved within mammalian precursors such as proenkephalin and prodynorphin, which encode multiple copies of the peptide sequences within a single prohormone ^62^. Although redundancy between peptides encoded in the same precursor has been described before ^54,55,57,61,63,64^, here we link changes in copy number to behavioral alterations. Such redundancy may sustain autocrine loops, amplify signaling, and preserve flexibility for context-specific functions. For instance, inhibitory autoreceptors may set a tonic baseline under resting conditions, whereas during stress or pathogen exposure, elevated *flp-1* expression could increase peptide availability, enabling more systemic release and activation of additional receptors. In line with this hypothesis, a previous study showed that *flp-1* expression increases following exposure to pathogenic bacteria, thereby enhancing signaling to RIM and RIC neurons, where the peptide drives locomotor avoidance ^29^.

We find that FLP-1/DMSR-7 signaling both self-regulates *flp-1* expression and controls the expression of other AVK-derived peptides, underscoring its role as homeostatic controller of neuropeptide expression in this hub neuron. While previous work on the FLP-1/DMSR-7 axis reported regulation only at the level of NLP-10 release, we additionally detect an effect on the expression of *flp-1*, amongst other neuropeptides. This discrepancy may be attributable to the use of different strategies, as our measurements rely on both mRNA quantification and a CRISPR-engineered insertion of a fluorescent reporter at the endogenous locus, which may reveal regulatory effects masked in the earlier study ^27^.

The molecular pathways acting downstream of FLP-1/DMSR-7 signaling in AVK, controlling neuropeptide expression, remain to be determined. A broad study of bHLH transcription factors identified four neuropeptide genes—*flp-1*, *nlp-49*, *nlp-50*, and *nlp-69*—as being regulated by HLH-15 in AVK ^47^. We did not find any change in expression of *nlp-69* upon removal of *flp-1*, whereas loss of *flp-1* has opposing effects on the expression of *nlp-49* and *nlp-50*. These results could point to additional transcription factors being involved in the regulation of AVK peptide expression by FLP-1. Additionally, bHLH transcription factors can function either as homodimers or heterodimers, therefore diversifying their gene-regulatory outcomes ^47^. Alongside HLH-15, the FAX-1 nuclear receptor and the UNC-42 homeobox gene have also been shown to regulate *flp-1* expression in AVK, with UNC-42 acting as a terminal selector for AVK identity throughout the animal’s life ^65^. Together, these findings raise the possibility that the FLP-1/DMSR-7 autocrine loop may act through one or more of these transcription factors to sustain neuropeptidergic expression, allowing context-dependent adjustments in peptide expression while preserving AVK neuronal identity.

Finally, not all AVK-expressed neuropeptides were regulated by FLP-1 signaling when looking at transcript levels in whole mount animal extracts, suggesting transcriptional selectivity rather than global scaling of AVK-expressed neuropeptides. Such selectivity may act to minimize crosstalk between signaling cascades while preserving AVK’s ability to control diverse behaviors. A recent example of hierarchical regulation within AVK is NLP-10, whose expression is independent of FLP-1, yet whose release is tightly controlled by FLP-1 to fine-tune the locomotor response ^27^. Future approaches combining cell-specific RNA-seq with peptide localization in dense-core vesicles could clarify how hub neurons are organized to enable such peptide-specific control. Beyond peptide expression, profiling of receptors, transcriptional regulators, and vesicle machinery may reveal how feedback reshapes neuronal molecular identity to sustain circuit homeostasis.

Taken together, our results suggest that AVK integrates opposing peptidergic feedback mechanisms, modulating the expression and release of AVK peptides, to control locomotor state. More broadly, they support the view that peptidergic hub neurons use autocrine feedback as a conserved strategy to keep peptidergic output in check while retaining adaptive capacity ^15,17,25,27,66^. By combining feedback mechanisms with functional redundancy between similar peptides, peptidergic hubs may achieve the balance of stability and flexibility that governs modulatory control.

### Limitations of the study

Although our findings establish an autocrine inhibitory role of DMSR-7 in regulating *flp-1* expression and release from AVK neurons, limitations remain. First, some of our analyses rely on whole-animal RNA measurements, which may obscure cell-specific transcriptional changes occurring within AVK and other neurons. Future single-cell transcriptomics could more precisely show variations at individual cellular level. Second, although we show that DMSR-7 acts cell-autonomously in AVK, we cannot completely rule out potential non-autonomous influences from other neurons—for example, those involved in PDF-1 signaling. Finally, we only performed behavioral assays on unstimulated animals; it remains to be tested whether the same mechanisms apply during stress conditions when a robust release of peptides is required.

## Structured Methods - Reagents and Tools Table

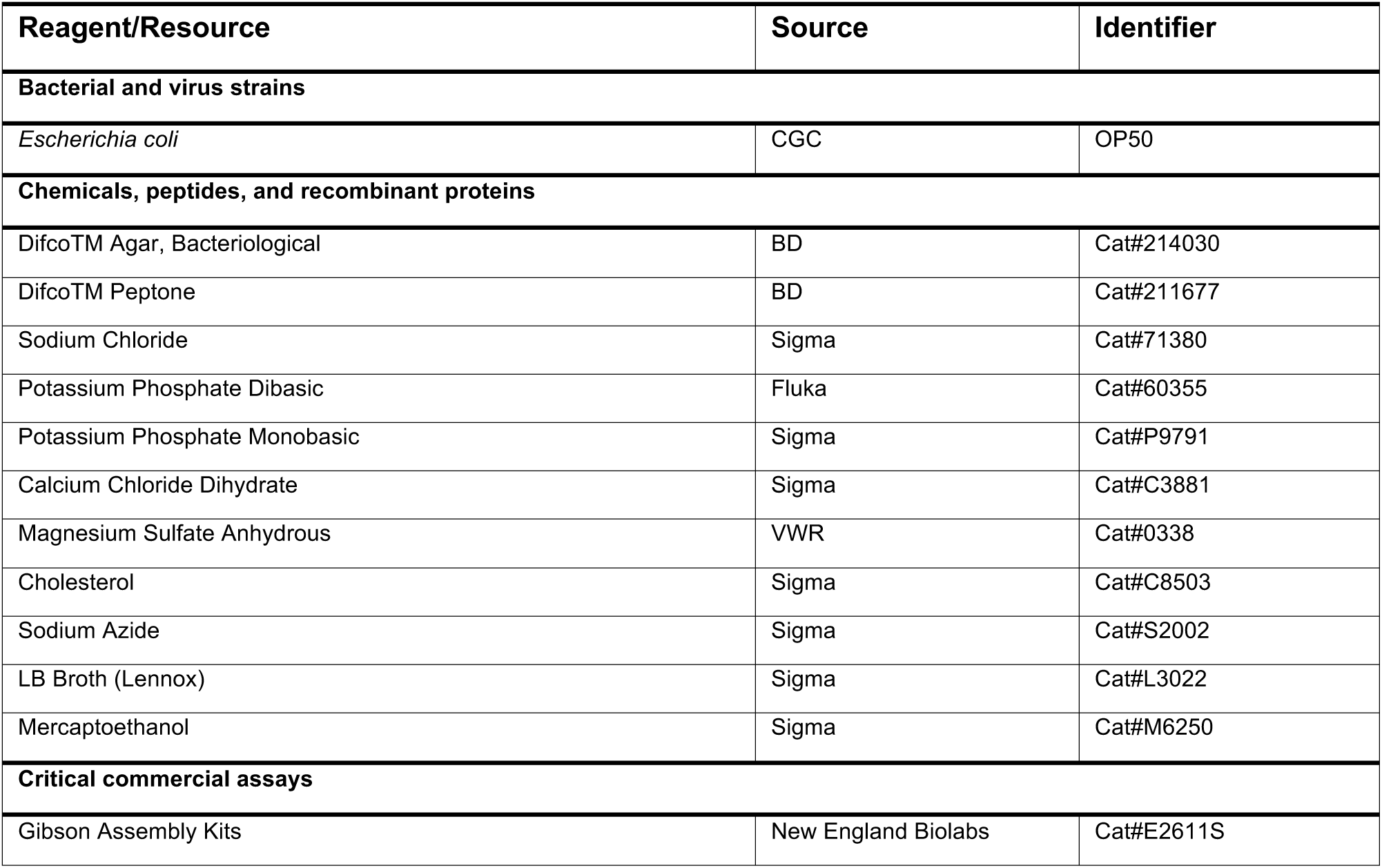
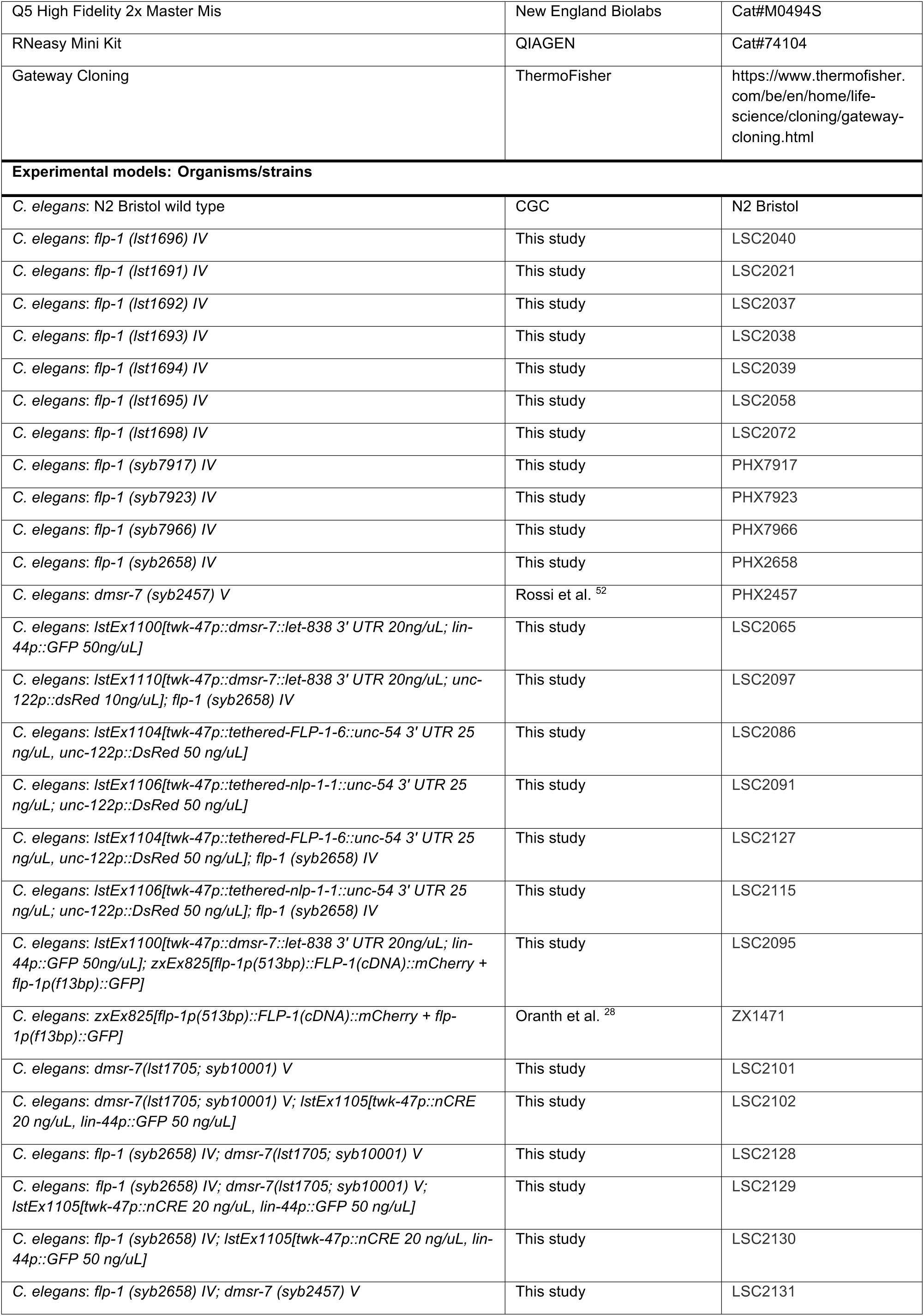
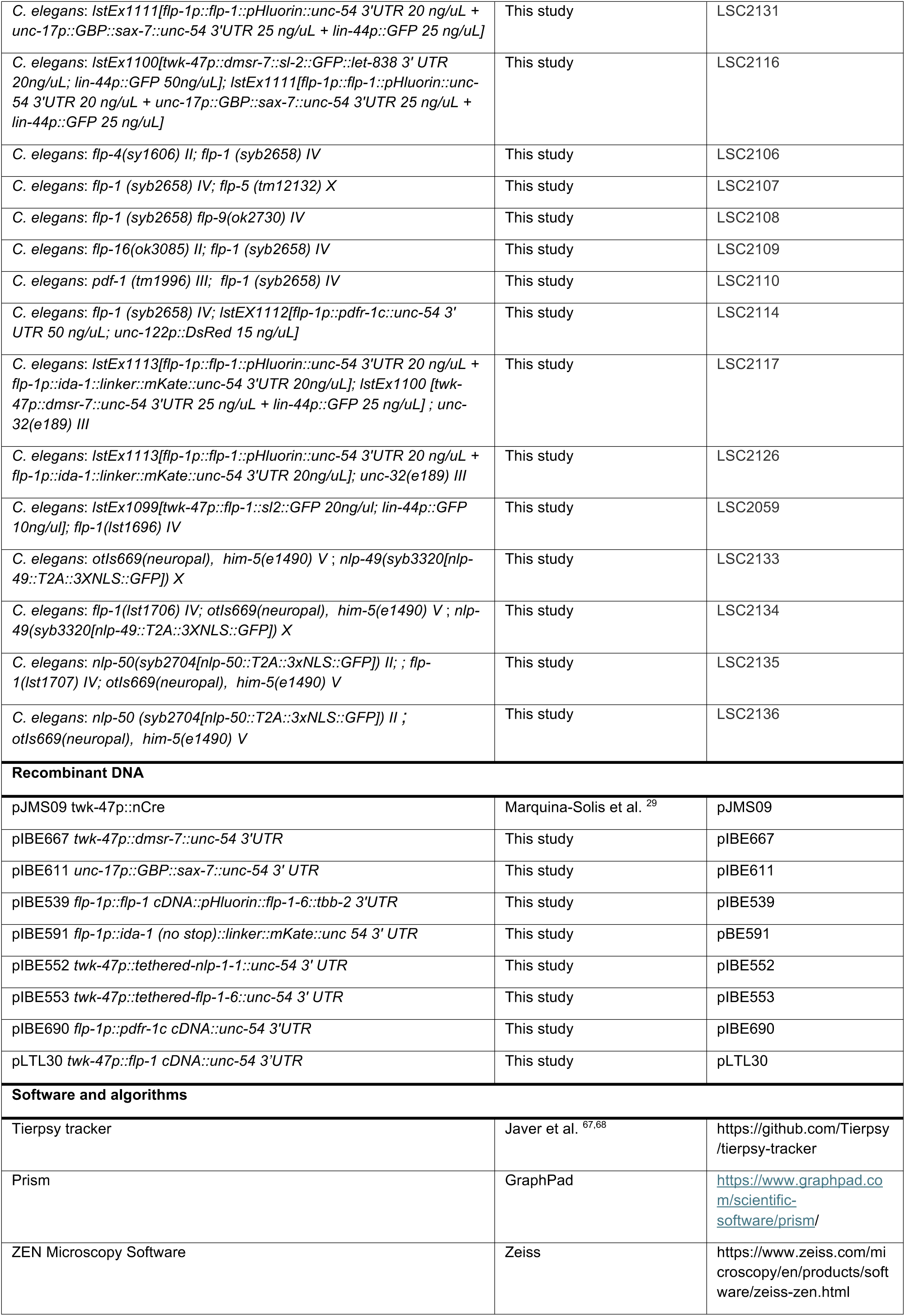
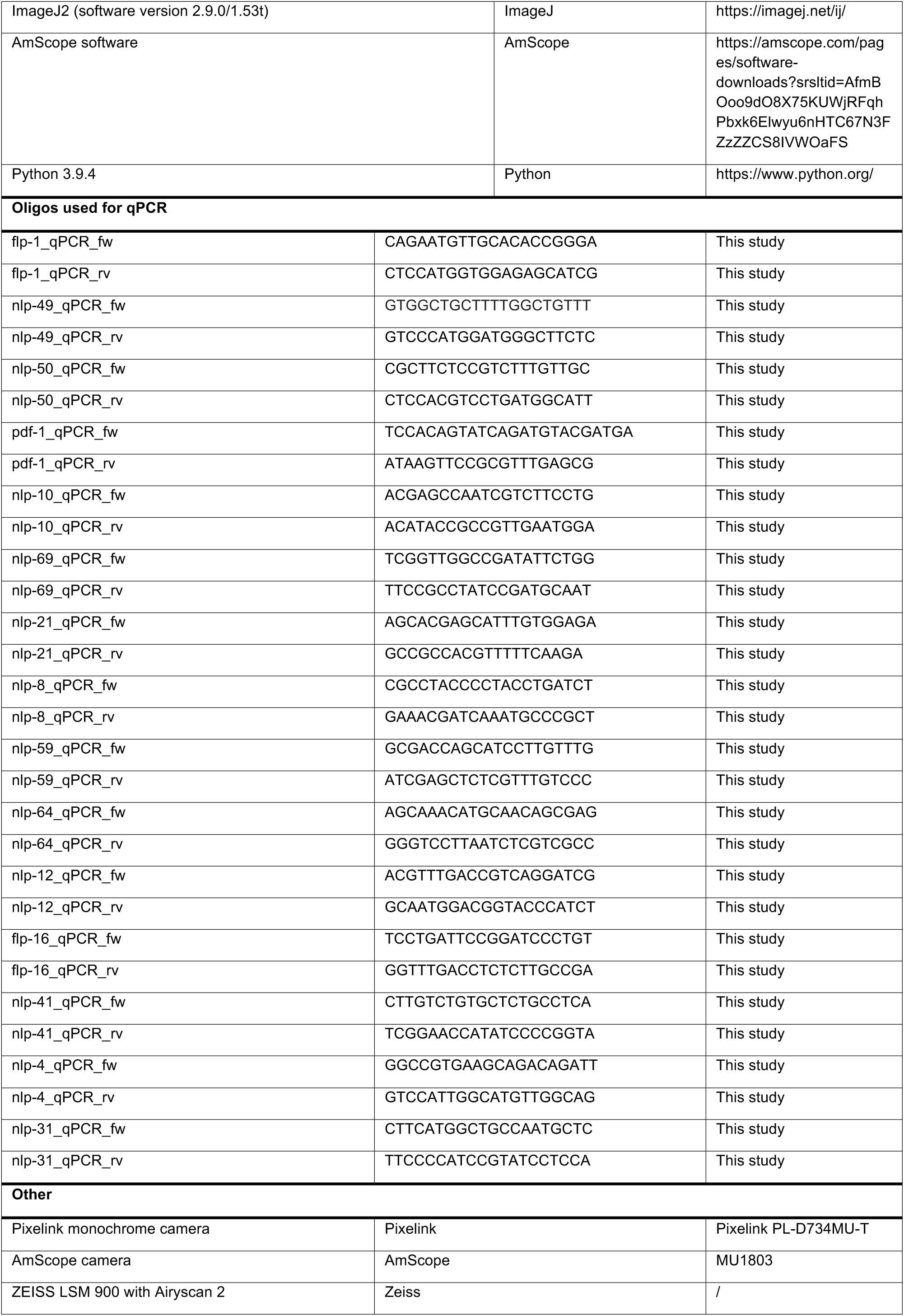

## Methods

### General maintenance

*C. elegans* was cultured at 20 °C, unless otherwise stated for specific assays, on nematode growth medium (NGM) and fed *E. coli* OP50 using standard methods ^69^. The wild-type N2 Bristol strain was obtained from the *Caenorhabditis* Genetics Center (University of Minnesota). The full list of strains used in this study can be found in the Experimental Models section of the Structured Methods - Reagents and Tools Table.

### Plasmid cloning and generation of transgenic strains

Clones used in this study were generated using Gibson Assembly (New England Biolabs) and MultiSite Gateway Three-Fragment (Invitrogen) cloning systems. *flp-1*(2865 bp), *twk-47* (250 bp) and *unc-17* (3291 bp) promoter regions were cloned from *C. elegans* mixed-stage gDNA using a Q5 high-fidelity polymerase (New England Biolabs). Plasmids with the *C. elegans* codon-optimized *pHluorin* and *GBP-sax-7* were a kind gift from Dr. Derek S. Sieburth (USC Keck School of Medicine, Los Angeles, USA). AVK-specific *twk-47p::Cre* plasmid was kindly provided by the lab of Cori Bargmann (Rockefeller University, New York, USA). The complete list of plasmids used in this study can be found in the Recombinant DNA section of the Structured Methods - Reagents and Tools Table. Transgenic animals were obtained by injection of young adult worms. F1 progenies were screened for transgene using a fluorescence microscope.

### CRISPR/Cas9 *flp-1* knockout and conditional *dmsr-7* knockout

Genetic editing was performed using a *dpy-10-*based co-CRISPR strategy ^24,70,71^. CRISPR RNAs (crRNAs) were chosen using CRISPOR ^72^. Young adults were injected with a mix of 0.25 µL Cas9 enzyme (15 ng/µL, Thermo Fisher), 0.26 µL tracrRNA (0.17 mol/l, Integrated DNA Technologies [IDT]), 0.24 µL *dpy-10* crRNA (0.6 nmol/μL, IDT), 0.20 µL *flp-1* crRNA1 (0.6 nmol/μL, IDT), 0.20 µL *flp-1* crRNA2 (0.6 nmol/μL, IDT), 2.2 µL *dpy-10* repair template (0.5 mg/ml, Merck), and 1.1 µL *flp-1* repair template (1 mg/ml, IDT). Progeny of the injected worms were screened by PCR for the desired gene edit. The genotype of the obtained mutants was verified through sequencing. The same protocol was adopted to generate the strain LSC2101 (conditional knockout of *dmsr-7*): the first LoxP site (ATAACTTCGTATAATGTATGCTATACGAAGTTAT) was inserted in the 5’ region of the gene, 100 bp upstream of the start codon ATG. A second insertion was performed at the end of the 3’ UTR of the gene, consisting of a LoxP site, followed by the sequence of *mKate::tbb-2 3’ UTR.* The complete sequence of the edit is reported in Supplementary file S1.

### Locomotion assay

One day prior to assaying, 10 L4-stage animals were transferred to fresh low peptone (0.130 g/L) NGM plates seeded with a drop of 10 µL OP50 and and maintained at 15°C. The following day, worms were recorded using a Pixelink PL-D734MU-T monochrome camera for 5 min at 15 fps using the manufacturer’s software (Pixelink Capture). Plates were positioned in the tracking system and allowed to acclimate for 2 minutes prior to recording. Videos were imported into Tierpsy Tracker using standard protocols ^67,68^. Animal segmentation and traces were manually inspected and corrected if necessary. From the resulting HDF5 files containing extracted features, mean speed and body curvature (at head, neck, midbody, hips, and tail segments) were calculated per plate, averaging across individuals. Data were imported in GraphPad Prism (version 8) for further analysis. One-way ANOVAs with post hoc Tukey’s tests were performed when comparing different strains. Significance is indicated in the figures as follows: * p < 0.05, ** p < 0.01, *** p < 0.001, **** p < 0.0001. Bar graphs show mean ± SEM. Raw data for locomotion assays included in this paper are available in Supplementary file S2.

### Swimming assay

Animals were bleach synchronized following the standard protocol ^73^. On the day of the assay, regular NGM containing 2.5 g/L peptone was freshly prepared. In each well of a 96-well Greiner plate with a clear, flat bottom, 100 μL of NGM was dispensed, dried, and covered with 100 μL of S-basal. Young adult worms were first transferred to an unseeded plate to remove any residual bacteria from their bodies. Following this step, 10 animals were transferred into each well and dispersed in S-basal solution. A 10-minute acclimation period was allowed before starting the video recording. Each well was recorded for 1 minute using an AmScope MU300 camera and AmScope software. Recordings were taken at 15 fps. Videos were blinded for analysis. Thrashing cycles were manually counted for each animal over a 30-second interval. The individual counts were then averaged per well. Statistical analysis was performed using GraphPad Prism (version 8). One-way ANOVAs with post hoc Dunnett’s tests were performed when comparing different strains. Significance is indicated in the figures as follows: * p < 0.05, ** p < 0.01, *** p < 0.001, **** p < 0.0001. Bar graphs show mean ± SEM. Raw data for swimming assays are available in Supplementary file S2.

### Confocal imaging

L4-stage fluorescent animals were handpicked under a fluorescence microscope the afternoon before imaging and allowed to grow overnight. Imaging slides were freshly prepared on the morning of the experiment using a 2% agar on glass slides. A 5 µL drop of M9 buffer was placed on the agar pad and the worms were dispersed in it. To immobilize the animals, a 5 µL drop of 1 µM sodium azide was added. A coverslip was then placed over the preparation to secure the sample.

Imaging was performed using a Zeiss LSM900 Airyscan 2 confocal microscope with 10× and 63× objectives and Immersol 518F (Zeiss) immersion solution. Z-stack images were acquired at 0.16 µm intervals and processed using the maximum intensity projection feature in Fiji/ImageJ2 (version 2.9.0/1.53t). Mean fluorescence intensity and distances between regions of interest (ROIs) were quantified using Fiji/ImageJ2. Data were imported into GraphPad Prism (version 8) for visualization, and statistical analysis was performed using one-way ANOVA followed by Tukey’s post hoc test. Significance is indicated in the figures as follows: * p < 0.05, ** p < 0.01, *** p < 0.001, **** p < 0.0001. Bar graphs show mean ± SEM. Raw data and statistics for quantified fluorescent images can be found in Supplementary file S2.

### RNA extraction and cDNA synthesis

Each strain was grown on 2.5 g/L peptone NGM plates and amplified via maternal bleaching following standard protocols. Synchronized L1 larvae in S-basal were counted and plated onto freshly seeded 90 mm NGM plates at a density of 5,000 worms per plate. Animals were grown at 15 °C until they reached the young adult stage. For each strain, four plates (totaling 20,000 worms) were washed with S-basal and collected into 15 mL tubes. Worms were washed three times to remove residual bacteria, then snap-frozen in liquid nitrogen for 1 minute and stored at −80 °C until RNA extraction.

On the day of extraction, worm pellets were resuspended in 600 μL of RLT buffer containing β-mercaptoethanol (100:1 concentration), according to the RNeasy Mini Kit (Qiagen, Cat. No. 74106) and transferred to MagNA Lyser Green Beads tubes. Samples were homogenized by shaking for 30 seconds at 6500 rpm using the MagNA Lyser, incubated at room temperature for 5 minutes, then transferred to 2 mL tubes and centrifuged at 4 °C for 3 minutes. The resulting supernatant was transferred to new tubes and mixed with 700 μL of ethanol. RNA was purified using RNeasy Mini Kit spin columns combined with DNase I (QIAGEN) treatment according to the manufacturer’s instructions. RNA concentration and purity were assessed using an Implen spectrophotometer. Samples with a 260/280 absorbance ratio of approximately 2.0 were considered pure.

For each strain, 1000 ng of total RNA was used to synthesize cDNA using the PrimeScript RT Reagent Kit (Takara, Cat. No. RR037A), resulting in cDNA at a final concentration of approximately 100 ng/μL.

### Quantitative real-time PCR

qRT-PCR was performed in 20 μL reaction mixtures, each containing 10 μL of Fast SYBR Green Master Mix (Thermo Fisher Scientific), 1.2 μL of a 10 μM gene-specific primer mix (see Table 1), 3.8 μL of Milli-Q water, and 5 μL of cDNA template. Reactions were performed in 96-well plates using the QuantStudio 3 Real-Time PCR System (Thermo Fisher Scientific). For each strain, at least 3 biological samples were tested in technical duplicates to ensure reproducibility.

Reference housekeeping genes *gpd-2* (glyceraldehyde 3-phosphate dehydrogenase), *rpb-12* (RNA polymerase II subunit), and *iscu-1* (involved in intracellular iron ion homeostasis) were included in the screening for normalization ^74,75^. The relative expression levels of *flp-1*, *flp-16*, *nlp-4*, *nlp-8*, *nlp-10*, *nlp-12*, *nlp-21*, *nlp-31*, *nlp-41*, *nlp-49*, *nlp-50*, *nlp-59*, *nlp-64*, *nlp-69*, and *pdf-1* were calculated using the ΔΔCt method with normalization to the geometric mean of the selected reference genes. Fold change (FC) values were plotted in GraphPad Prism (version 8) and differenced across strains analyzed using one-way ANOVA with Tukey’s post hoc test. Raw data can be found in Supplementary file S2.

## Data availability

All data generated in this study is reported in this paper. Further information and material should be directed to the <u>lead contact</u> Liesbet Temmerman

This paper does not report original code.

Any additional information required to reanalyze the data reported in this paper is available from the <u>lead contact</u> upon request.

## Author contributions

Conceptualization, L.G, I.B., and L.T.; methodology, L.G, I.B., and L.T.; Investigation, L.G.; writing—review & editing, L.G, I.B., and L.T.; funding acquisition, I.B. and L.T.; supervision, I.B. and L.T.

## Acknowledgments

We thank Prof. Cori Bargmann for kindly sharing the plasmid pJMS09 (*twk-47p::Cre*), Prof. Derek Sieburth for sharing plasmids encoding *pHluorin* and *sax-7::GBP* and Prof. Alexander Gottschalk for sharing the strain ZX1471. We thank Dr. Ichiro Aoki and Prof. Alexander Gottschalk for valuable feedback on the manuscript. We thank Amanda Kieswetter, Elke Vandewyer, Keertana Venkatesh, Dr. Jan Watteyne, Julian Matytchak and Len De Paep for technical support, and are grateful to the *Caenorhabditis* Genetics Center (CGC) [supported by the National Institutes of Health Office of Research Infrastructure Programs (P40 OD010440)] for providing strains.

## Supplemental information

**Supplementary File S1.** CRISPR engineered strains produced in this paper

**Supplementary File S2.** Raw data and statistics presented in this paper

**Figure S1.**
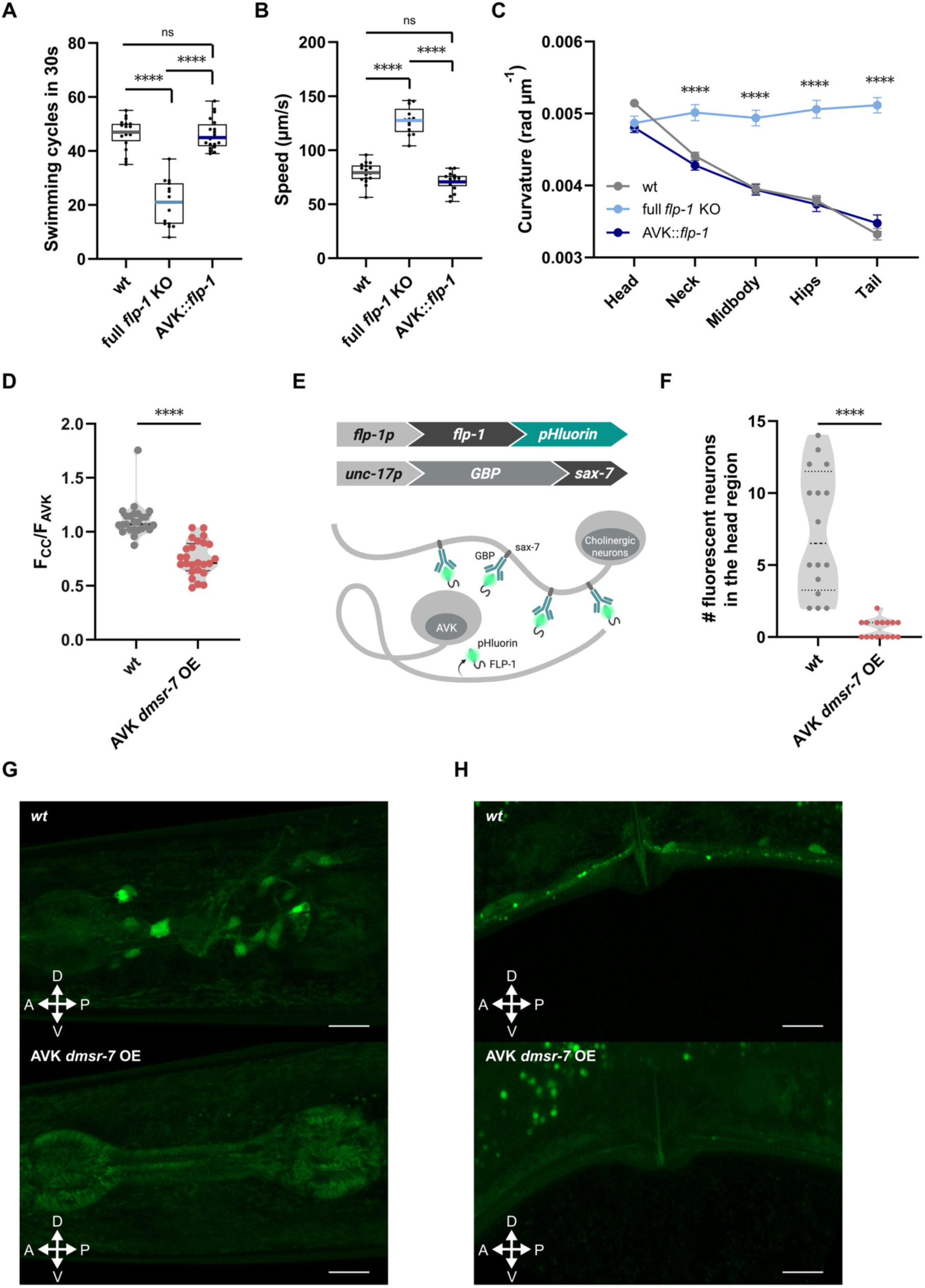
Quantification and visualization of FLP-1 distribution in wild-type and *dmsr-7*-overexpressing animals. **(A)** Swimming cycles of *flp-1* mutant animals and mutants rescued in AVK. **(B and C)** Speed and curvature for full *flp-1* mutants and rescued animals in AVK neurons. (**D)** Mean fluorescence intensity of FLP-1::mKate in coelomocytes, normalized to AVK fluorescence, quantified in wild-type and *dmsr-7*-overexpressing animals. (**E)** Schematic of the biochemical strategy used to visualize FLP-1 distribution. FLP-1 peptide, expressed under its promoter and tagged with pHluorin, interacts with membrane-anchored (sax-7) GFP Binding Protein (GBP) expressed in cholinergic neurons (under *unc-17p*). **(F)** Number of fluorescent neurons in the head of wild type and animals overexpressing *dmsr-7* in AVK. **(G and H)** Confocal images of cholinergic neurons tethering released FLP-1::pHluorin in head (D) and vulva (E) regions of wild-type and *dmsr-7*-overexpressing animals. Scale bar = 15 µm. Data represent mean ± SEM from **(A to C)** ≥ 14 individual plates with 10 animals in each, and **(D and F)** ≥ 15 animals per genotype. Statistical analysis performed using one-way ANOVA with post hoc Dunnett’s tests **(A),** Tukey test in **(B and C)** and t-test for **(D and F)**; ****P < 0.0001.

**Figure S2.**
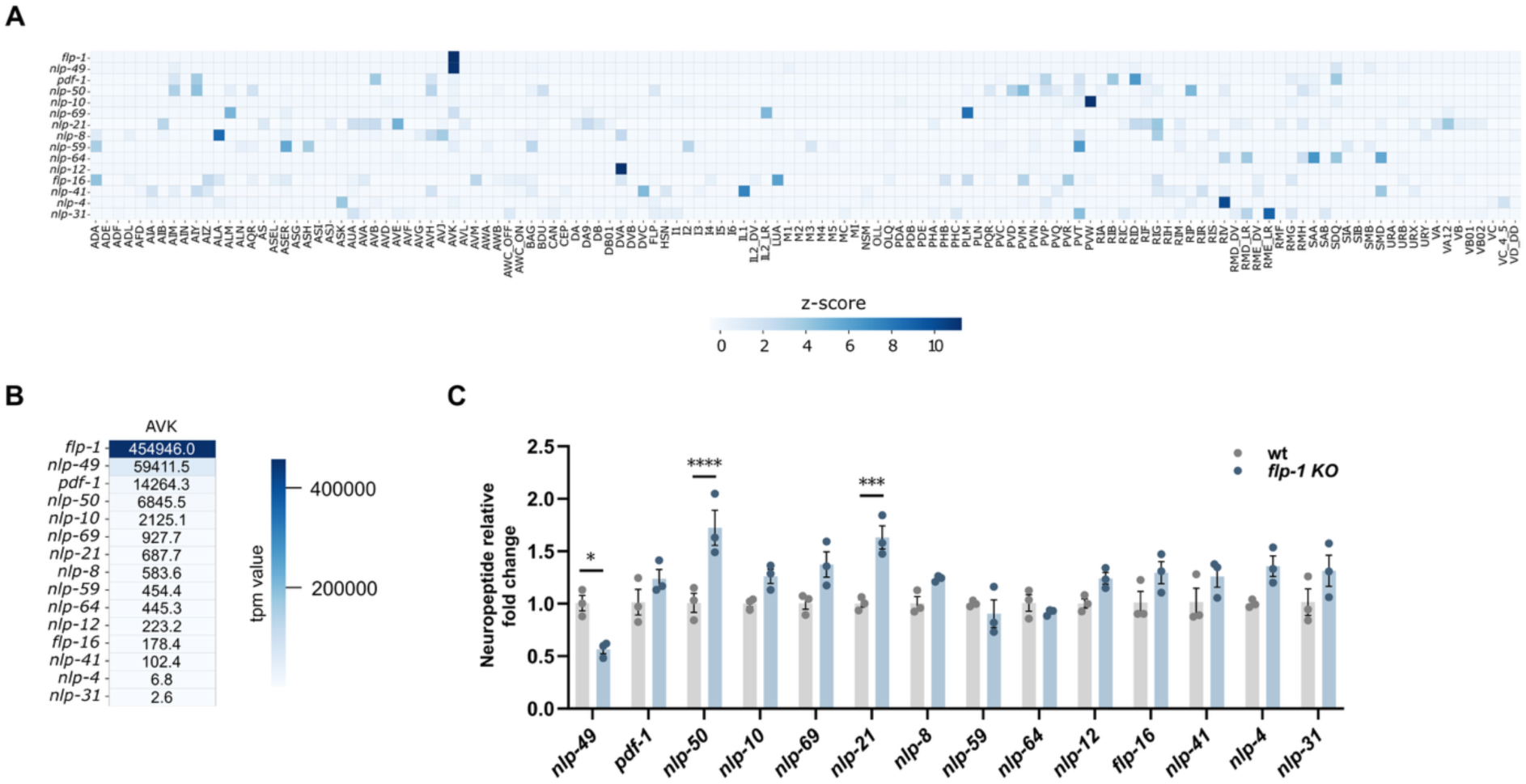
Expression and regulation of neuropeptides in AVK neurons. **(A)** Z-score values of neuropeptides expressed in AVK neurons across the entire nervous system of L4-stage animals ^34^. **(B)** Transcripts per million (TPM) expression values of neuropeptides detected in AVK neurons. **(C)** Relative fold change in neuropeptide expression in wild-type versus *flp-1* knockout animals from whole-mount animal mRNA extrats. Data represent mean ± SEM from N = 3 independent assays. Statistical analysis was performed using one-way ANOVA followed by post hoc Tukey correction; ****P < 0.0001 ***P < 0.001, *P < 0.05.

**Figure S3.**
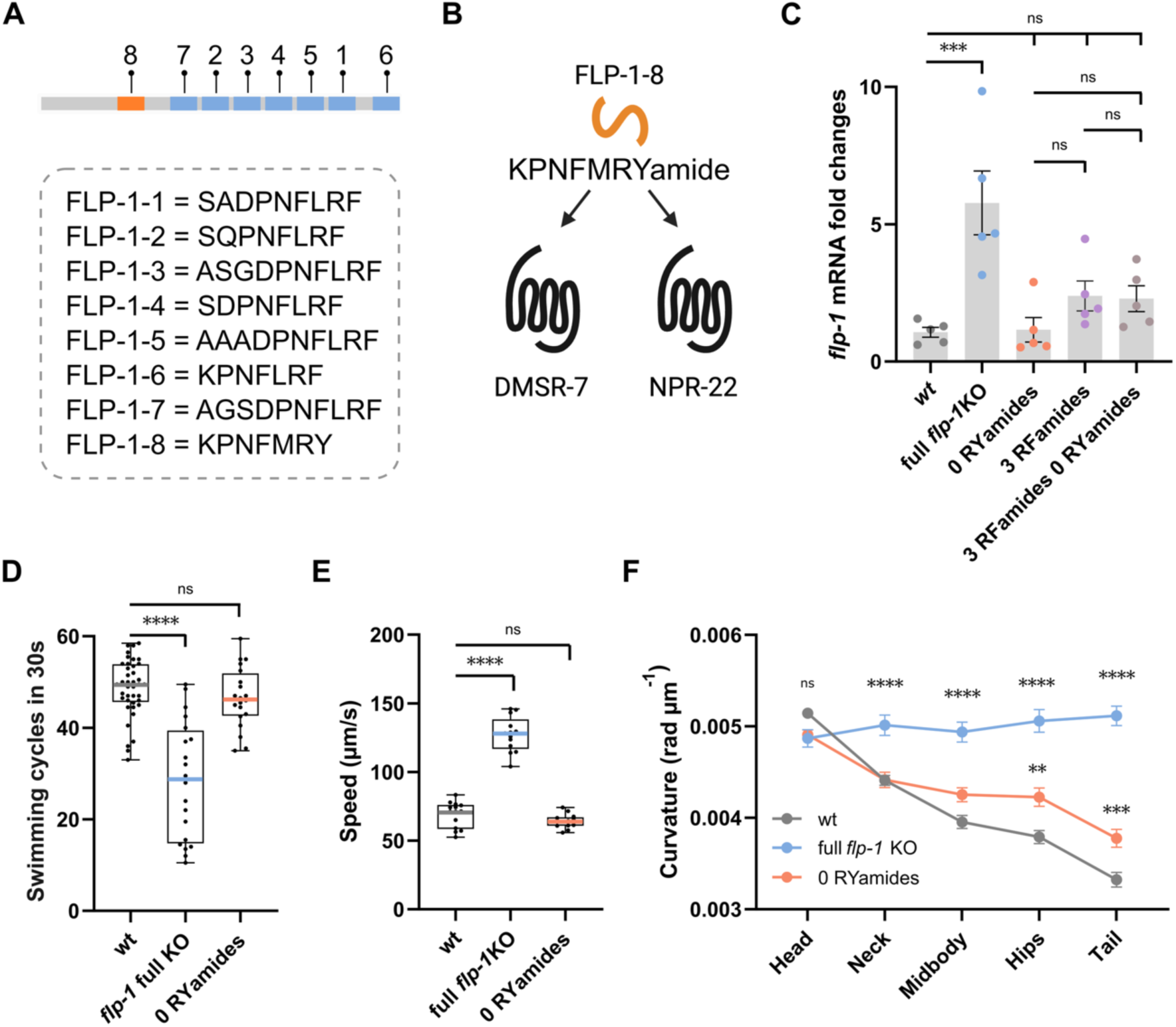
FLP-1 RYamide signaling doesn’t regulate locomotor behavior. **(A)** Sequences for individual FLP-1 peptides retrieved from Beets et al., 2023 ^20^. **(B)** Schematic representation of the FLP-1-8 RYamide peptide and its two cognate receptors identified through in vitro assays ^20^. **(C)** Relative *flp-1* mRNA expression levels in wild-type, *flp-1* knockout, 3RFamide peptide knockout, RYamide peptide knockout, and 3RFamide + RYamide peptide knockout animals. **(D)** Quantification of mean swimming cycles performed over 30 seconds in wild-type, *flp-1*, and RYamide peptide knockout animals. **(E and F)** Locomotor parameters of animals crawling on food: **(E)** mean speed and **(F)** mean body curvature across five body segments. Data in **(D to F)** represent mean ± SEM from ≥ 14 independent assays. Data in **(C)** represent mean ± SEM from 3 independent assays. Statistical analysis was conducted using one-way ANOVA followed by post hoc Tukey test **(C, E and F)** and Dunnett’s tests **(D)**; ****P < 0.0001, ***P < 0.001, **P < 0.01.

